# Alveolar regeneration following viral infection is independent of tuft cells

**DOI:** 10.1101/2022.03.11.483948

**Authors:** Huachao Huang, Ming Jiang, Yihan Zhang, Jana Biermann, Johannes C. Melms, Jennifer A. Danielsson, Yinshan Fang, Ying Yang, Li Qiang, Jia Liu, Yiwu Zhou, Manli Wang, Zhihong Hu, Timothy C. Wang, Anjali Saqi, Jie Sun, Ichiro Matsumoto, Wellington Cardoso, Charles W. Emala, Jian Zhu, Benjamin Izar, Hongmei Mou, Jianwen Que

## Abstract

Severe injuries following viral infection cause lung epithelial destruction with the presence of ectopic basal progenitor cells (EBCs), although the exact function of EBCs remains controversial. We and others previously showed the presence of ectopic tuft cells in the disrupted alveolar region following severe influenza infection. Here, we further revealed that the ectopic tuft cells are derived from EBCs. This process is amplified by Wnt signaling inhibition but suppressed by Notch inhibition. Further analysis revealed that *p63-CreER* labeled population de novo arising during regeneration includes alveolar epithelial cells when Tamoxifen was administrated after viral infection. The generation of the *p63-CreER* labeled alveolar cells is independent of tuft cells, demonstrating segregated differentiation paths of EBCs in lung repair. EBCs and ectopic tuft cells can also be found in the lung parenchyma post SARS-CoV-2 infection, suggesting a similar response to severe injuries in humans.

## Introduction

While the deadly “Spanish” influenza pandemic killed approximately 50 million people worldwide during 1918-1920, seasonal influenza (flu) continued to be deadly claiming 291,000 to 646,000 lives each year (CDC, 2017). Moreover, the COVID-19 pandemic caused by SARS-CoV-2 virus has already claimed over 5.9 million lives and infected over 432 million people thus far (JHU, 2022). Post-mortem examination of H1N1 influenza-infected lungs revealed extensive tissue remodeling accompanied by the presence of ectopic cytokeratin-5 positive (KRT5^+^) basal cells in the lung parenchyma (Xi et al., 2017). These ectopic basal cells (EBCs) are also present in the mouse parenchyma following infection with modified H1N1 influenza PR8 virus or treatment with a high dose of bleomycin (Kanegai et al., 2016; Vaughan et al., 2015; Xi et al., 2017; Yuan et al., 2019; Zacharias et al., 2018; Zuo et al., 2015). Initial studies suggest that these EBCs contribute to alveolar regeneration (Zuo et al., 2015). However, studies from other groups suggest that EBCs do not meaningfully become alveolar epithelial cells, but rather provide structural supports to prevent the lung from collapsing (Basil et al., 2020; Kanegai et al., 2016; Vaughan et al., 2015). These seemingly contradictory results necessitate a better understanding of EBCs.

Tuft cells are a minor cell population critical for chemosensory and relaying immune signals in multiple organs including the intestine and trachea (Howitt et al., 2016; Montoro et al., 2018; von Moltke et al., 2016). In the intestine tuft cells serve as immune sentinels during parasitic infection (Gerbe et al., 2016; Howitt et al., 2016; McGinty et al., 2020b; von Moltke et al., 2016). Responding to helminth infection, tuft cells release the alarmin interleukin (IL)-25 which activates type 2 innate lymphoid cells (ILC2s) and their secretion of IL-13, initiating type 2 immune response to eliminate parasitic infection (Gerbe et al., 2016; Howitt et al., 2016; von Moltke et al., 2016). Tuft cells were initially identified in the rat trachea over six decades ago (Rhodin and Dalhamn, 1956). However, we just have begun to appreciate their functions in the respiratory system (Bankova et al., 2018; Ualiyeva et al., 2020). Recent single cell RNA sequencing confirmed the presence of tuft cells in the mouse trachea where they express several canonical genes, such as *Alox5ap, Pou2f3*, and *Gfi1b* (Montoro et al., 2018). Following repeated allergen challenges, tuft cells expand and amplify the immune reactions (Bankova et al., 2018). Intriguingly, tuft cells were found ectopically present in the lung parenchyma, co-localized with EBCs following PR8 viral infection (Rane et al., 2019). More recently, we reported that ectopic tuft cells were also present in the parenchyma of COVID-19 lungs, and ablation of tuft cells dampens macrophage infiltration at the acute phase following viral infection in a mouse model (Melms et al., 2021). However, the role of tuft cells in lung regeneration following viral infection remains undetermined. It is also unknown what cells generate these tuft cells and what signaling pathways control the derivation.

In this study, we show that EBCs serve as the cell of origin for the ectopic tuft cells present in the parenchyma during viral infection. Upon screening multiple signaling pathways we identified that Notch inhibition blocks tuft cell derivation, while Wnt inhibition significantly enhances tuft cell differentiation from EBCs. We then used multiple mouse models to demonstrate that *p63-CreER* labeled subpopulations of alveolar type I and 2 (AT1/AT2) cells during regeneration regardless of the presence of tuft cells in the parenchyma.

## Results

### Tuft cells expanded in response to challenges with influenza, bleomycin or naphthalene

Tuft cells are rarely present in the proximal airways of normal adult mice (Montoro et al., 2018). However, they were present in the distal airways at birth when tuft cells were first detected (Figure 1A and 1B). Tuft cells continued to be present throughout the airways when examined at postnatal (P)10 and P20 but were no longer detected in the terminal bronchiole at P56 (Figure 1A and 1B). Notably, tuft cells reappeared in the terminal airways approximately 7 days post infection (dpi) with PR8 virus (Figure 1C). The number of tuft cells also significantly increased in the large airways at 30 dpi (4.9 ± 1.1 Vs 25.7 ± 3.3 per 1000 epithelial cells) (Figure 1C). As previously described, tuft cells were present in the EBC area (Melms et al., 2021; Rane et al., 2019). Further analysis revealed that tuft cells were first detected in the EBC area at approximately 15dpi and peaked at 21dpi (Figure E1A). The numbers of tuft cells also expanded in the large airways following naphthalene (13.4 ± 1.1 per 1000 epithelial cells) or bleomycin challenge (8.6 ± 0.8 per 1000 epithelial cells) (Figure 1D). In addition, tuft cells were ectopically present within EBCs 28 days following challenge with a high dose of bleomycin (Figure 1D). These findings suggest that tuft cell expansion in the airways is a general response to lung injuries, and that severe injuries induce ectopic tuft cells in the parenchyma.

**Figure 1.**
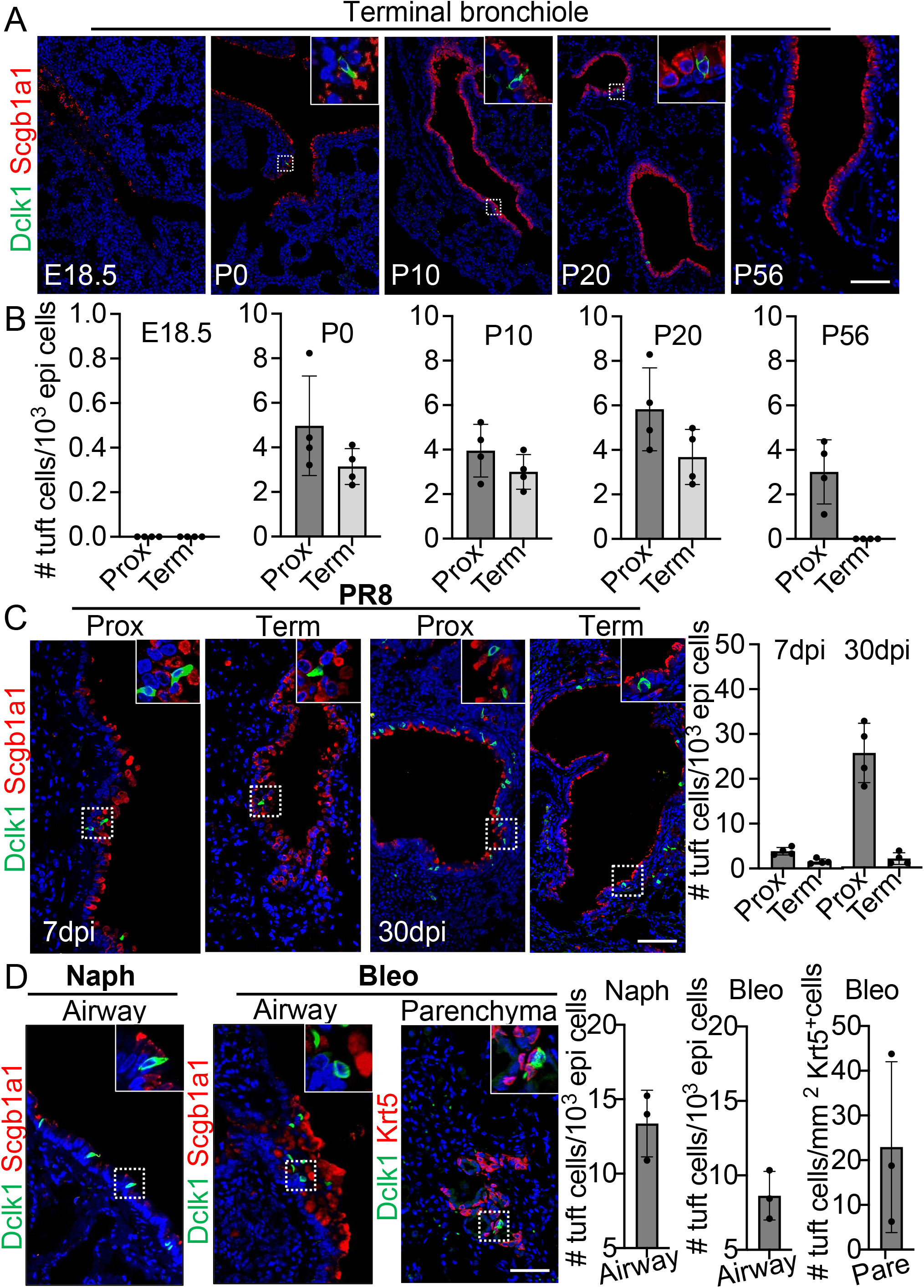
Tuft cells in homeostasis and their expansion in response to severe injuries. (*A*) Tuft cells are present in the distal airways at P0, P10, P20 but not P56. n=4 for each group. (*B*) Quantification of tuft cells from panel (A). (*C*) Tuft cells expand in the large airway and are ectopically present in the terminal airways following viral infection. (*D*) Tuft cells expand in the airways following naphthalene and bleomycin challenge and are ectopically present in parenchyma following bleomycin challenge. Scale bars, 50 μm.

### The ectopic tuft cells in the parenchyma did not proliferate or give rise to other cell lineages

Previous studies suggest that Dclk1^+^ cells serve as progenitor cells in the small intestine and pancreas (May et al., 2008; May et al., 2009; Westphalen et al., 2016). We asked whether the ectopic tuft cells in the lung parenchyma also possess progenitor potential. We therefore performed lineage tracing with a knock-in *Pou2f3-CreER* mouse line (McGinty et al., 2020b). Tuft cells (Dclk1^+^) in the large airways were specifically labeled with the lineage tracing maker (tdT^+^) upon three Tamoxifen (Tmx) injections into *Pou2f3-CreER;R26tdT* mice (Figure 2A). Notably, ~92% tdT^+^ cells expressed Dclk1 in the trachea, whereas all tdT^+^ cells expressed Dclk1 in the large intrapulmonary airways (Figures 2A and Supplement file S1B). We then analyzed the mice challenged with PR8 virus. Tmx was continuously given from 14 to 27 dpi and the lungs were analyzed at 60 dpi (Figure 2B). The majority (~93%) of the ectopic tuft cells were lineage labeled in the lung parenchyma of *Pou2f3-CreER;R26tdT* mice (Figure 2B). These labeled tuft cells solitarily distributed within EBCs (Figure 2B), indicating that they had not undergone expansion. Consistently, none of the tuft cells were positive for the proliferation marker Ki67 (Figure 2C). To further test whether tuft cells give rise to other cell lineages, immunostaining with various cell type markers was performed, including Scgb1a1 (club cells), FoxJ1 (ciliated cells) and Clca3 (goblet cells). The results showed that tuft cells did not contribute to other cell types following a chasing period up to 60 days (Figure 2D). These data suggest that tuft cells in the lung unlikely serve as progenitor cells.

**Figure 2.**
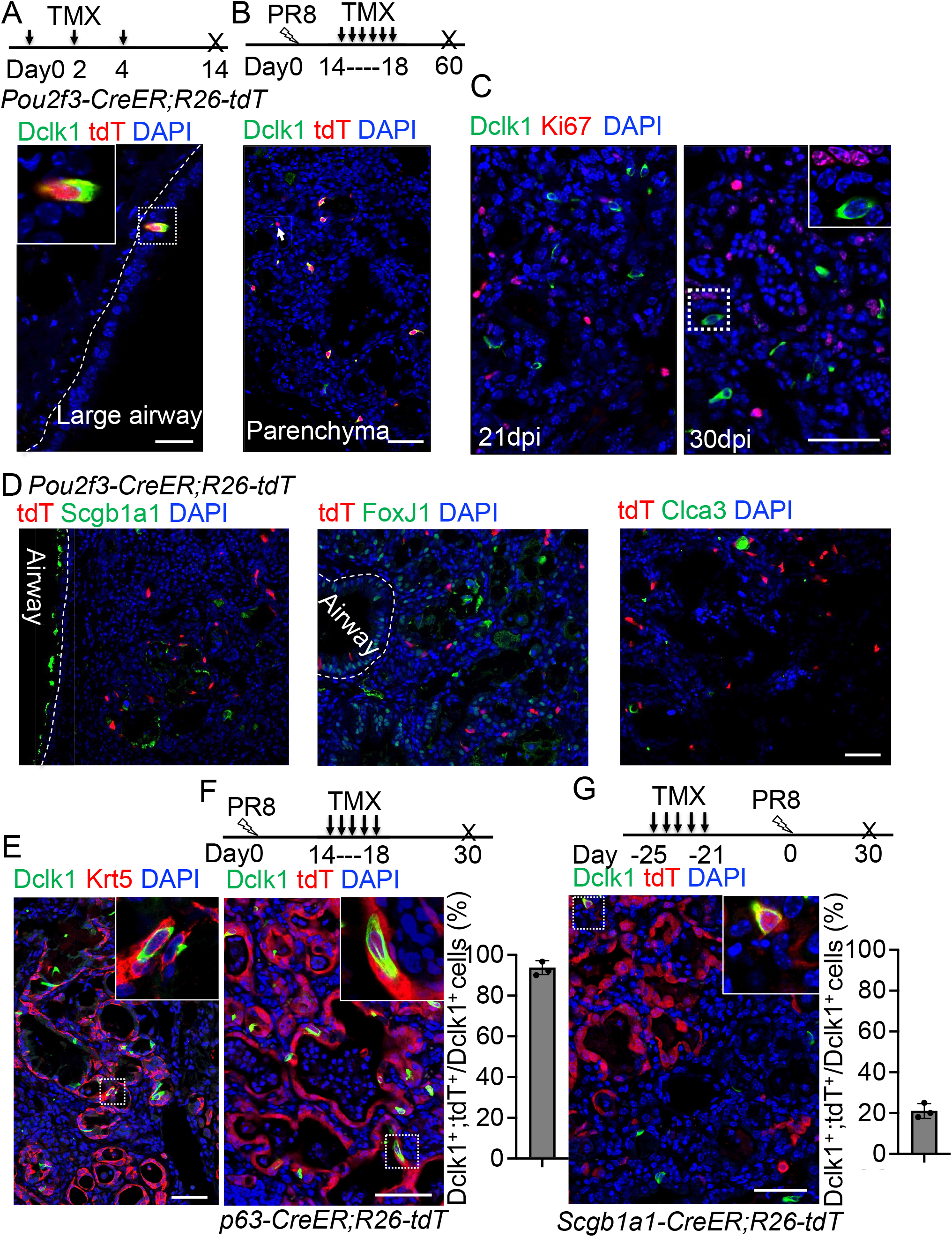
Tuft cells in the parenchyma are derived from ectopic basal cells (EBCs) following H1N1 PR8 viral infection. (*A*) *Pou2f3-CreER;R26-tdT* mouse line specifically labels Dclk1^+^ tuft cells in the large airways at homeostasis. (*B*) Linage tracing of Pou2f3^+^ cells in the parenchyma. Arrow indicates a tdT^+^Dclk1^-^ cell. (*C*) Tuft cells do not proliferate when examined at days 21 and 30 post infection. (*D*) Lineage labeled tuft cells do not contribute to other cell lineages. (*E*) ~5 % of tuft cells within EBCs coexpress Krt5 and Dclk1. (*F*) Lineage tracing confirms that tuft cells are derived from EBCs in the lung parenchyma of *p63-CreER;R26tdT* mice following PR8 infection. (*G*) Lineage labeled cells contribute to ~21% ectopic tuft cells in the parenchyma of *Scgb1a1-CreER;R26-tdT* mice. Data represent mean ± s.e.m. \**p*<0.05, \*\**p*<0.01; statistical analysis by unpaired two-tailed Student’s *t*-test. Scale bars, 20 μm (A) and 50 μm (B to G).

### The ectopic tuft cells in the parenchyma originated from EBCs

Basal cells serve as the origin of tuft cells in the mouse trachea (Montoro et al., 2018). By contrast, lineage-negative epithelial stem/progenitor cells (LNEPs) were considered as the origin of tuft cells in the lung parenchyma following viral infection (Rane et al., 2019). While characterizing ectopic tuft cells in the parenchyma, we noticed approximately 5% tuft cells co-expressed Dclk1 and the basal cell marker Krt5 (Figure 2E), suggesting that these tuft cells are in a transitioning state from EBCs towards tuft cells. To test this hypothesis, *p63-CreERT2;R26tdT* mice were infected with PR8 virus. Given that tuft cells were initially detected at around 15dpi, we injected Tmx daily from 14 dpi to 18 dpi (Figure 2F). The majority of tuft cells within EBCs were labeled with tdT (Figure 2F), suggesting that EBCs serve as the cell origin for the ectopic tuft cells. EBCs can be derived from multiple cell sources including a subpopulation of Scgb1a1^+^ club cells (Yang et al., 2018). To test whether EBCs derived from the club cell subpopulation give rise to tuft cells, we lineage labeled club cells before exposing *Scgb1a1-CreER;R26tdT* mice to PR8 virus. 21.0% ± 2.0 % tuft cells within EBCs are labeled with tdT when examined at 30 dpi (Figure 2G), suggesting that club cell-derived EBCs generate a portion of tuft cells in the lung parenchyma upon viral infection.

### Inhibition of Wnt and Notch signaling has opposite effects on the differentiation of EBCs into tuft cells

We next sought to identify the signaling pathways that can influence tuft cell differentiation from EBCs. Lineage-labeled pod cells were isolated and purified from the lungs of *p63-CreER; R26tdT* mice following viral infection and were expanded using the protocol for culturing basal cells (Mou et al., 2016) (Supplement file S2A). The expanded EBCs maintained the lineage tag (tdT^+^) and expressed the basal cell markers p63 and Krt5 (Supplement file S2B). In addition, they also expressed the respiratory protein markers Nkx2.1 and Foxa2 (Supplement file S2B). We then used the expanded EBCs to assess the impact of major signaling pathways which have been implicated in the determination of cell fate during lung development. The tested pathways include Tgfß/Bmp, Yap, Shh, Wnt, Notch, IL-6 and IL-4/IL-13 (Ahdieh et al., 2001; Barkauskas et al., 2013a; Ikonomou et al., 2020; Lee et al., 2014b; Li et al., 2017; Nabhan et al., 2018; Pardo-Saganta et al., 2015; Tadokoro et al., 2014; Vaughan et al., 2015; Yuan et al., 2019; Zacharias et al., 2018). We observed that treatment with IL-4 or IL-13 apparently promoted tuft cell derivation from EBCs in air-liquid interface (ALI) culture (data not shown), in agreement with the previous reports of IL-4Rα-dependent tuft cell expansion from the intestinal crypts (Gerbe et al., 2016; von Moltke et al., 2016). Notably, treatment of the WNT signaling activator CHIR9902 blocked the derivation of tuft cells by 35.3% ± 3.1%. Conversely, treatment with the Wnt inhibitor IWR-1 promoted tuft cell differentiation by 32.4% ± 6.4 % (Supplement file S2C). Inhibition of WNT signaling with the Porcupine inhibitor Wnt-C59 also led to increased tuft cell differentiation of airway basal cells isolated from *Dclk1-GFP* mice (Figure 3A). Consistently, daily injection of Wnt-C59 induced abundant tuft cells in both airways and lung parenchyma following viral infection (Figure 3B and Supplement file S2D). By contrast, Notch inhibition with the γ-secretase inhibitor Dibenzazepine (DBZ) reduced tuft cell differentiation from EBCs in ALI culture (Figure 3C). Daily DBZ injection also decreased the number of tuft cells in the parenchyma by 63.6% ± 1.5% following viral infection (Figure 3D). Together these findings suggest that inhibition of the WNT signaling pathway promotes while Notch inhibition blocks the differentiation of EBCs into tuft cells.

**Figure 3.**
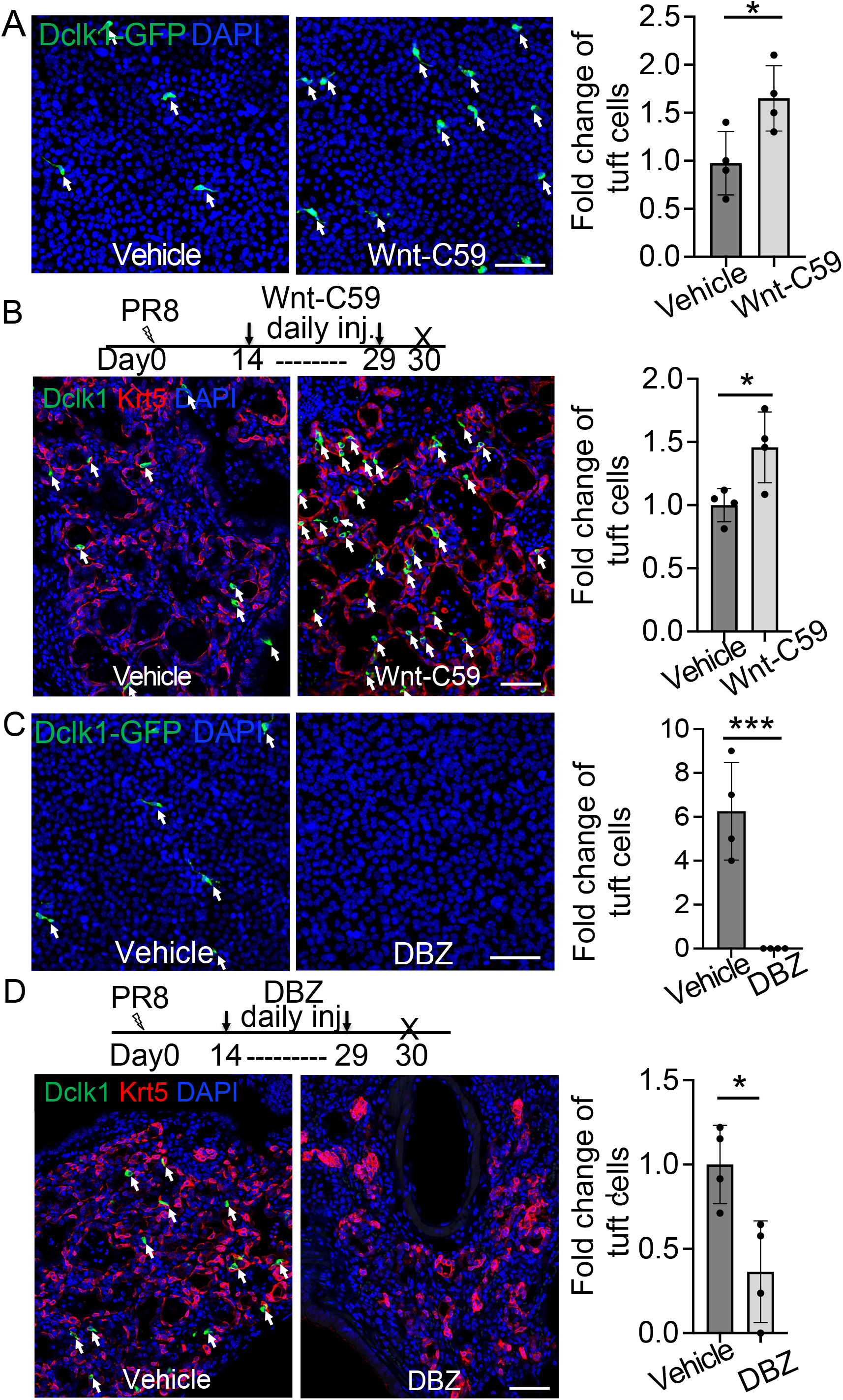
Wnt inhibition promotes EBC differentiation into tuft cells while Notch inhibition has an opposite effect. (*A*) Treatment with the WNT signaling inhibitor Wnt-C59 promotes tuft cell differentiation of *Dclk1-GFP* basal cells in ALI culture. (*B*) Wnt-C59 treatment increases the number of tuft cells in the parenchyma following viral infection. (*C*) Treatment with the Notch signaling inhibitor DBZ completely blocks tuft cell differentiation of *Dclk1-GFP* basal cells in ALI culture. (*D*) Daily injection of DBZ following PR8 infection dramatically reduces tuft cell derivation in the parenchyma. n=4 per group. Data represent mean ± s.e.m. \**p*<0.05, \*\*\**p*<0.001; statistical analysis by unpaired two-tailed Student’s *t*-test. Scale bars, 50 μm.

### Enhanced generation of *p63-CreER* lineage labeled alveolar epithelium in *Trpm5^-/-^* but not *Pou2f3^-/-^* mutants

Controversy remains regarding the contribution of EBCs to alveolar regeneration (Vaughan et al., 2015; Zuo et al., 2015). Given tuft cell ablation reduces macrophage infiltration in PR8-infected mouse lungs (Melms et al., 2021), we asked whether loss of tuft cells improved alveolar regeneration. The first mouse model we examined was *Trpm5* null mutant which demonstrates ~80% reduction of tuft cells in the intestine (Howitt et al., 2016). However, the number of tuft cells were not significantly reduced following viral infection in the lung parenchyma of *p63-CreER;Trpm5^+/-^;R26tdT* mice as compared to control mice (*p63-CreER;Trpm5^+/-^;R26tdT*) (p>0.05, Supplement file S3A). Furthermore, we did not observe an apparent reduction in the size of Krt5^+^ EBC clones in the lung parenchyma (Supplement file S3B). Notably, 19.4 % ± 4.2% *p63-CreER* lineage-labeled cells expressed the AT2 cell marker SftpC but not Krt5 as compared to the control (7.9% ± 1.4%) when Tmx was injected after viral infection (Figures 4A and Supplement file S3C). Lineage labeled AT2 cells were further confirmed by Abca3, another mature AT2 cell marker (Supplement file S3D). In addition, lineage-labeled cells also include AT1 epithelium (Hopx^+^, T1a^+^) which was increased in the parenchyma of *p63-CreER;Trpm5^-/-^;R26tdT* mutants (6.5% ± 0.9% Vs 2.5% ± 0.8% at 60 dpi) (Figure 4B and Supplement file S3E). To assess whether *Trpm5^-/-^* mice exhibited improved pulmonary mechanics after viral infection, we assessed airway resistance when challenged with increasing concentrations of methacholine. Total respiratory system resistance (Rrs) and central airway resistance (Rn) were significantly attenuated in *Trpm5^-/-^* mice (Figure 4C).

**Figure 4.**
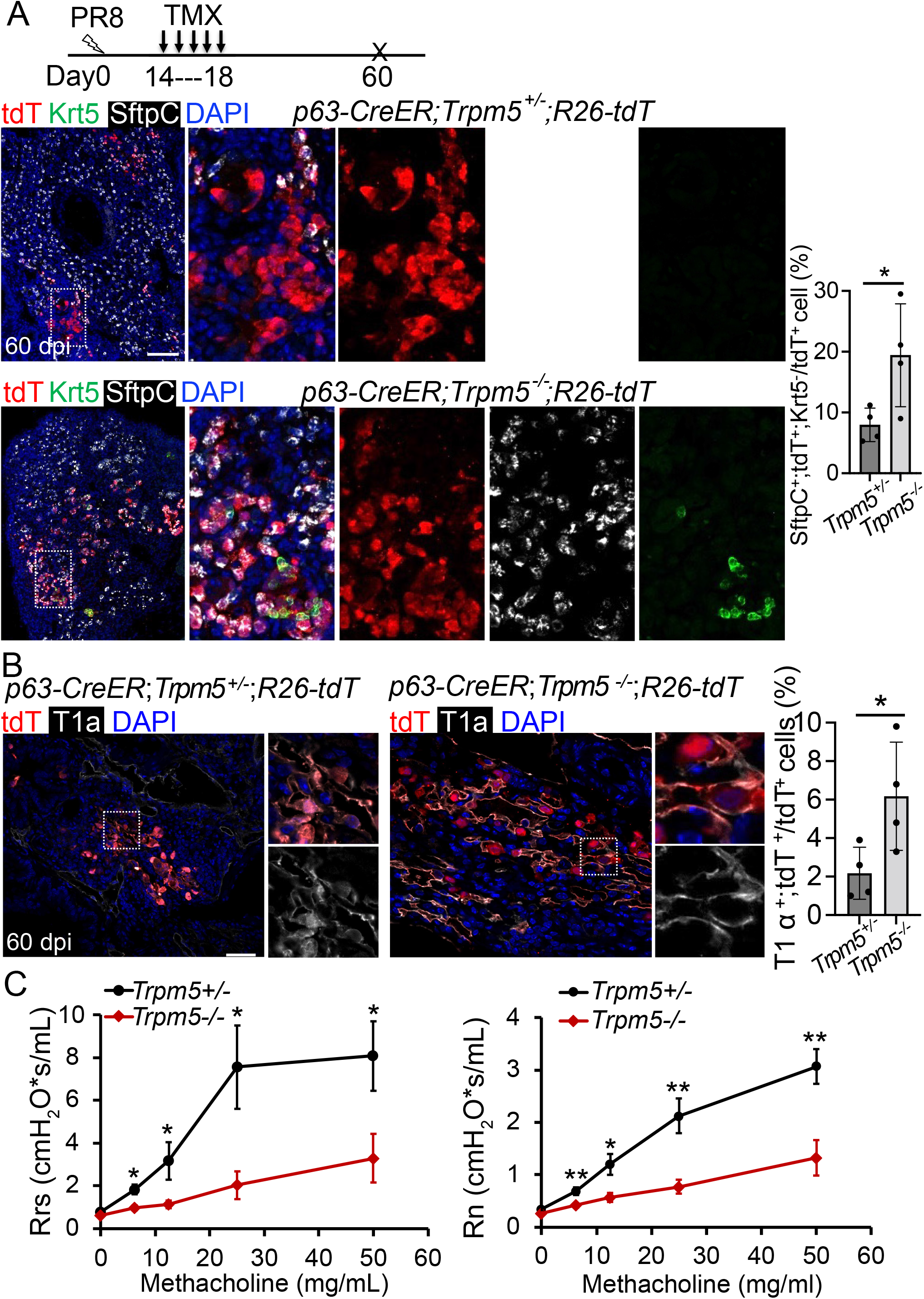
Increased generation of *p63-CreER* labeled alveolar epithelium in *Trpm5* null mutants following PR8 infection. (*A*) *Trpm5* deletion leads to the increased presence of tdT^+^SftpC^+^Krt5^-^cells in the parenchyma at 60 dpi. (*B*) Increased tdT-labeled AT1 cells in the mutant lungs as compared to controls at 60 dpi. (*C*) Whole lung airway resistance improves in *Trpm5^-/-^* mice following viral infection (left panel). Central airway resistance is also reduced following viral infection (right panel). n=7 for WT group, n=9 for *Trpm5^-/-^* group. Data represent mean ± s.e.m. \**p*<0.05, \*\**p*<0.01; statistical analysis by unpaired two-tailed Student’s *t*-test. Scale bars, 100 μm (A), 20 μm (B).

We next examined the contribution of *p63-CreER* lineage-labeled cells to lung regeneration in another mutant (*Pou2f3^-/-^*) which has no tuft cells (Matsumoto et al., 2011). We subjected *p63-CreER;Pou2f3^-/-^;R26tdT* mutants and controls (*p63-CreER;Pou2f3^+/-^;R26tdT*) to viral infection followed by continuous Tmx injection from 14 to 18 dpi. While prominent bronchiolization occurred in the parenchyma with the extensive presence of Krt5^+^ EBCs (Figure 5A), approximately 8% of the lineage-labeled cells became Sftpc^+^ Krt5^-^ AT2 cells in the controls (Figure 5B). However, we did not detect improved contribution of lineage-labeled alveolar epithelium in the mutants as compared to the controls (p>0.05) (Figure 5B), suggesting that loss of ectopic tuft cells has no impact on alveolar regeneration.

**Figure 5.**
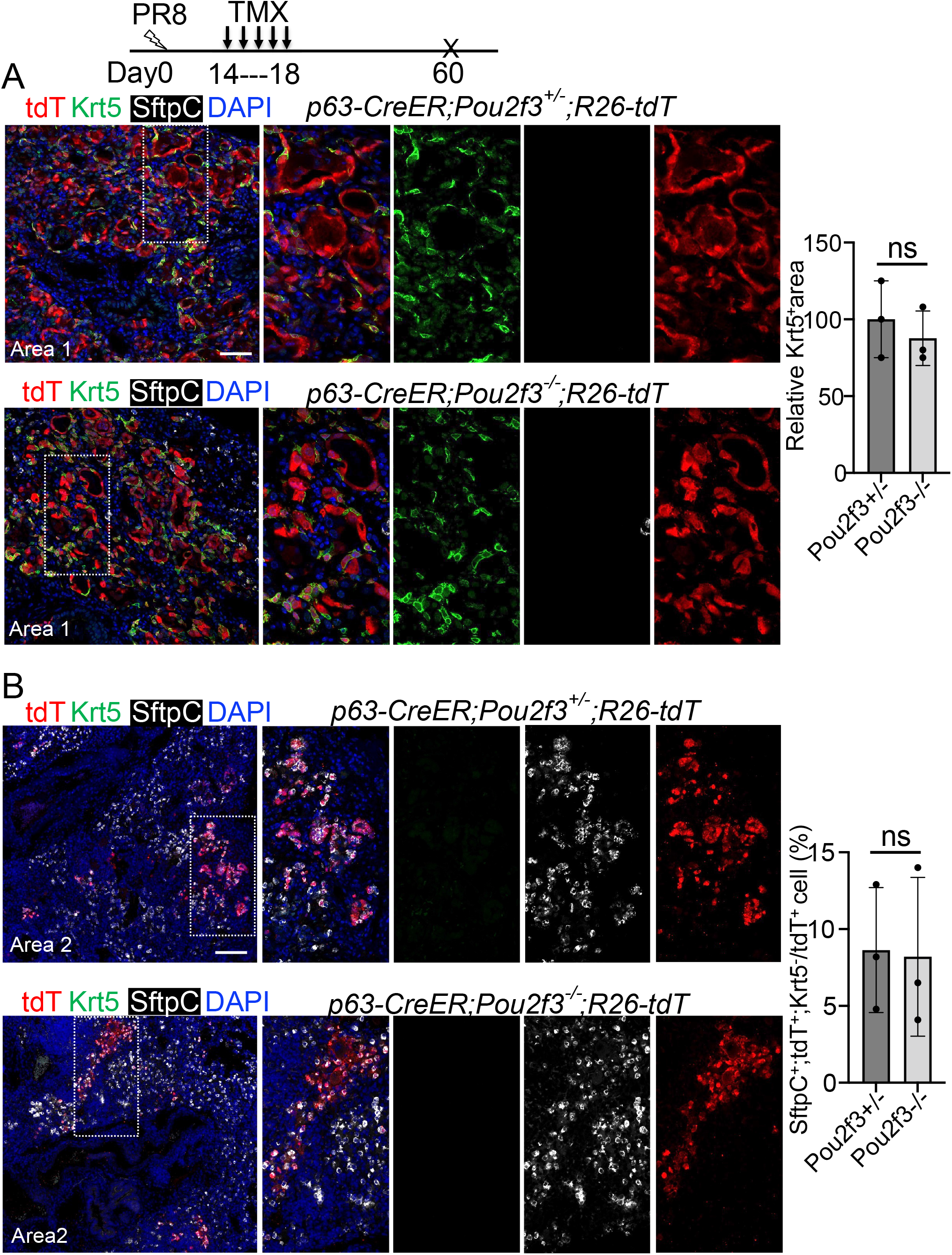
A complete loss of tuft cell does not increase the generation of *p63-CreER* labeled alveolar cells following PR8 infection. (*A*) Extensive Krt5^+^SftpC^-^ EBCs are present in the parenchyma of control (*p63-CreER;Pou2f3^+/-^;R26-tdT*) and mutant lungs (*p63-CreER;Pou2f3^-/-^;R26-tdT*) when examined at 60 dpi following PR8 infection. (*B*) Representative areas show *p63-CreER* labeled AT2 cells in both control and mutant lungs. ns: not significant; statistical analysis by unpaired two-tailed Student’s *t*-test. Scale bar, 50 μm (A), 100 μm (B).

### COVID-19 lungs contain EBCs which likely give rise to tuft cells

SARS-CoV-2 infection causes catastrophic damage to the lungs (Huang et al., 2020), and we recently reported expansion of tuft cell in COVID-19 lungs (Melms et al., 2021). SARS-CoV-2 entry into host cells requires the serine protease Transmembrane Serine Protease 2 (TMPRSS2) and Angiotensin-converting enzyme 2 (ACE2), which are present in club cells, ciliated cells and AT2 cells (Hoffmann et al., 2020; Lukassen et al., 2020). Consistently, club and ciliated cells were almost completely ablated (Fang et al., 2020), thus exposing the underlying basal cells in the affected intrapulmonary airways (Figure 6A). A significant number of the dispositioned basal cells remained proliferative while lodged in the alveoli, presumably initiating EBCs (Figure 6B-C). Tuft cells were occasionally present in the established EBCs in addition to the airways (Figure 6D). Similar to the EBC-derived tuft cells in influenza-infected mice, the ectopic tuft cells in the parenchyma of COVID-19 lungs co-expressed KRT5 and POU2F3 (Figure 6D), suggesting a similar differentiation scheme. By contrast, tuft cells in the airways did not express KRT5 (Figure 6E).

**Figure 6.**
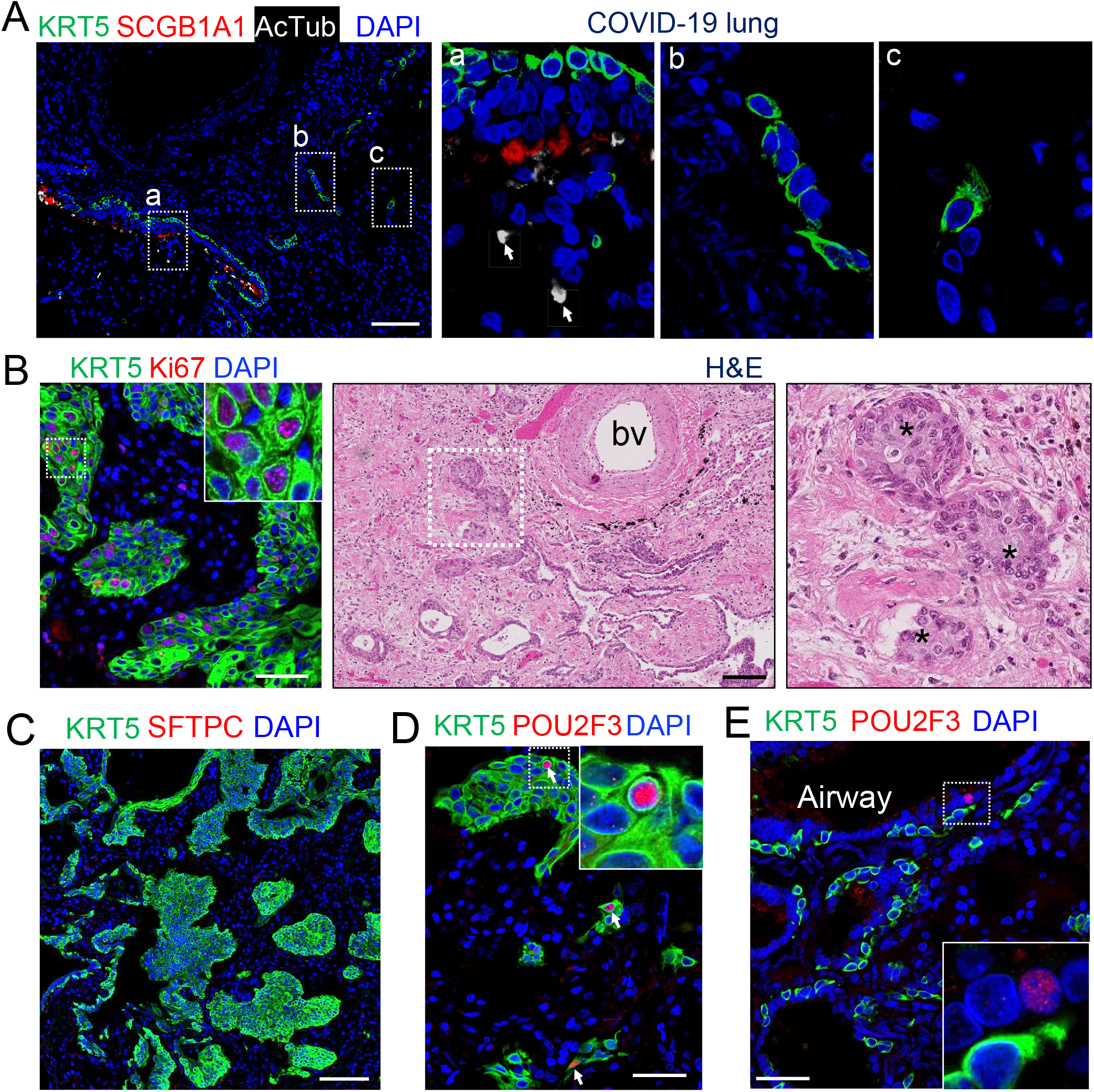
EBCs likely give rise to tuft cells in the parenchyma of COVID-19 lungs. (*A*) SARS-CoV-2 infection causes the loss of club and ciliated cells (arrows in a), exposing the underlying basal cells (b) in the small airways. Note the detached basal cells (c). (*B*) EBCs proliferate in the parenchyma of COVID-19 lungs. H&E staining shows the presence of EBC clusters in COVID-19 lungs. (*C*) Representative clusters of EBCs are present in COVID-19 lungs. (*D*) Tuft cells within EBCs express both KRT5 and the tuft cell marker POU2F3. (*E*) Solitary tuft cells without KRT5 expression are present in the airways of COVID-19 lung. Abbreviation: bv, blood vessel. Scale bars, 100 μm (A, B and C) and 50 μm (D and E).

## Discussion

Tuft cells are observed in the large airways at homeostasis in adults. Here, we showed that tuft cells are present in both large and terminal airways during early postnatal development. In response to severe injuries, tuft cells expanded in the airways. Additionally, tuft cells were ectopically present in the parenchyma of severely injured lungs, and lineage tracing confirmed that they were derived from EBCs. Inhibition of Wnt signaling promoted while Notch inhibition blocked EBC differentiation into tuft cells. Moreover, enhanced generation of the *p63-CreER* labeled alveolar cells was observed in *Trpm5^-/-^* but not *Pou2f3^-/-^* mutant. Finally, our study suggests that ectopic tuft cells in the parenchyma of COVID-19 lungs are derived from EBCs.

Tuft cells were detected throughout the airways including the terminal airways at around the neonatal stage, and they were no longer present in the terminal airways when examined at P56. Upon viral infection, tuft cells expanded in large airways and re-appeared in terminal airways, suggesting re-activation of developmental signaling during injury-repair. Our pilot screen of signaling inhibitors/activators demonstrated that Wnt blockage resulted in significantly increased differentiation of EBCs towards tuft cells. In consistence, treatment of the Wnt inhibitor Wnt-C59 also led to significant expansion of tuft cells in the airways and parenchyma. Along this line, influenza infection has been shown to cause downregulation of Wnt signaling in mouse lungs (Hancock et al., 2018). By contrast, we found that Notch inhibition significantly suppressed tuft cell expansion in the virus-infected lung parenchyma. Additionally, IL-4 and IL-13 treatments increased the derivation of tuft cells from EBCs, which is in contrast to the findings from the co-submitted manuscript where deletion of *IL-4* had no impact on tuft cell differentiation. This could be due to the extra IL-4/IL-13 we supplied to the cell culture, which may not be present in an in vivo setting during viral infection. Re-expression of the transcription factor Sox9 has been observed during the repair of the airway epithelium following naphthalene challenge (Jiang et al., 2021). Interestingly, Sox9 is expressed by tuft cells as shown by the accompanied manuscript. It will be interesting in future experiments to determine whether Sox9 is required to promote the generation of tuft cells.

The contribution of EBCs to alveolar regeneration remains controversial (Kanegai et al., 2016; Vaughan et al., 2015; Yuan et al., 2019; Zuo et al., 2015). Initial studies suggest that EBCs (also known as pods) serve as stem cells and are essential for lung regeneration (Zuo et al., 2015). However, other groups demonstrated that EBCs provide structural support to prevent the collapse of severely damaged lung without meaningful contribution to regeneration (Kanegai et al., 2016; Vaughan et al., 2015). Here, we used lineage tracing to demonstrate that *p63-CreER* labeled cells include AT1 and AT2 cells regardless of the mouse lines with/without tuft cells (*p63-CreER;R26dtT;Trpm5^+/-^*, *p63-CreER;R26dtT;Trpm5^-/-^, p63-CreER;R26dtT;Pou2f3^+/-^, p63-CreER;R26dtT;Pou2f3^-/-^*). EBCs have been reported to derive from multiple sources including Krt5^+^ basal cells (Vaughan et al., 2015; Zuo et al., 2015). In those studies, and the study accompanying this manuscript, Tmx was administrated prior to viral infection, presumably labeling preexisting basal cells with the *Krt5-CreER* mouse line. Lineage labeled cells did not contribute meaningfully to the regenerated alveolar cells. By contrast, we performed Tmx injection after viral infection and observed significant contribution of *p63-CreER* lineage labeled cells to the alveolar epithelium. It is possible that we labeled bona fide newly generated cell populations that expressed p63, although the cell origin for these cells remains to be determined. In addition to the preexisting basal cells, subpopulations of EBCs can be derived from Scgb1a1^+^ cells following viral infection (Yang et al., 2018). More recently, AT2 cells were also found to generate basal cells in vitro (Kathiriya et al., 2022). It is also possible that our strategy may label these club cells or AT2 cell-derived EBC subpopulations during the transition. In addition, at the early stage of mouse lung development (embryonic day 10.5), lineage-labeled p63^+^ progenitor cells give rise to AT1 and AT2 cells (Yang et al., 2018). Therefore it is also possible that subpopulations of AT1/AT2 cells regain the transcription program of fetal lung progenitors and transiently express p63 prior to becoming alveolar cells (Yang et al., 2018).

We noticed an increased contribution of *p63-CreER* labeled cells to alveolar cells in *Trpm5^-/-^* but not *Pou2f3^-/-^* mutants which lack tuft cells, suggesting that lung regeneration is independent of the presence of ectopic tuft cells in the parenchyma. Trpm5 is a calcium-activated channel protein that induces depolarization in response to increased intracellular calcium (Prawitt et al., 2003). It is also expressed in B lymphocytes in addition to tuft cells (Sakaguchi et al., 2020). We observed decreased accumulation of B lymphocytes in the lungs of *Trpm5^-/-^* mutants following viral infection (unpublished data, H.H. and J.Q.). It will be interesting in the future to determine whether reduced B lymphocytes facilitate lung regeneration.

In summary, we demonstrated that tuft cells are present in the airways at the early postnatal stages and later are restricted to the large airways. In response to severe injuries tuft cells expand in the airways and are ectopically present in the parenchyma where EBCs serve as their progenitor cells. Moreover, we identified *p63-CreER* labeled alveolar epithelial cells arising during lung regeneration independent of tuft cells.

## Methods and Materials

### Mouse Models

*p63-CreER* (Lee et al., 2014a), *Scgb1a1-CreER* (Rawlins et al., 2009), *Pou2f3-CreER* (McGinty et al., 2020a), *Trpm5-GFP* (Clapp et al., 2006), *Trpm5^-/-^* (Damak et al., 2006), and *Pou2f3^-/-^* (Matsumoto et al., 2011) mouse strains were previously described. *B6.Cg-Gt(ROSA)26Sor^tm14(CAG-tdTomato)Hze^/J(R26-tdT*) mouse strain was purchased from The Jackson Laboratory (Stock #007914). All mice were maintained on a C57BL/6 and 129SvEv mixed background and housed in the mouse facility at Columbia University according to institutional guidelines. 8-12 weeks old animals of both sexes were used in equal proportions. All animal studies used a minimum of three mice per group. Mouse studies were approved by Columbia University Medical Center (CUMC) Institutional Animal Care and Use Committees (IACUC).

### Administration of tamoxifen

Tamoxifen was dissolved in sunflower seed oil to 20 mg/mL as stock solution. For lineage analysis and genetic targeting, *Pou2f3-CreER* mouse strain were administered 2 mg tamoxifen by oral gavage at days 0, 2, and 4 for control or 1 mg tamoxifen by oral gavage at days 14, 17, 20, 23, 25, 27 post viral infection. *p63-CreER* mice were administered with tamoxifen at a dose of 0.25 mg/g bodyweight by daily oral gavage for a total five doses as indicated. For *Scgb1a1-CreER* mice, a period of ten or twenty-one days as indicated was used to wash out the residual tamoxifen before any further treatments.

### Injury models (influenza, bleomycin and naphthalene)

For influenza virus infection, 260 plaque forming units (pfu) of influenza A/Puerto Rico/8/1934 H1N1 (PR8) virus were dissolved in 40 μl of RPMI medium and then pipetted onto the nostrils of anesthetized mice, whereupon mice aspirated the fluid directly into their lungs. Post procedure, mice were weighed weekly and monitored for mortality rate. For bleomycin injury, mice were anesthetized and intratracheally instilled with 4 unit/kg body weight of bleomycin hydrochloride. For naphthalene treatment, naphthalene solution (25mg/ml) was freshly prepared before the procedure by dissolving naphthalene in sunflower seed oil. A single dose of naphthalene was delivered by intraperitoneal injection at 275 mg/kg body weight. For all procedures listed above, the administration of the same volumes of vehicle (PRMI medium or saline or sunflower seed oil) was used as control.

### Mouse EBC isolation, culture and differentiation

*p63-CreER; R26tdT* mice were infected with PR8 influenza virus and were administered with tamoxifen intraperitoneally as indicated. At 18dpi mouse peripheral lungs were dissected and dissociated according to the protocol as previously described (Barkauskas et al., 2013b; Rock et al., 2011). tdT^+^ cells were sorted by FACS and cultured using the protocol as previously reported (Mou et al., 2016), and the protocol for inducing the differentiation of basal-like cells was previously described (Feldman et al., 2019; Mou et al., 2016). Air-liquid interface (ALI) culture was used to test the effects of the major signaling pathway inhibitors on the differentiation of EBCs towards tuft and mucous cell lineages. Moreover, tracheal basal cells isolated from *Trpm5-GFP* mice were cultured and tested for drug effects in ALI. The tested inhibitors include the BMP signaling inhibitor Dorsomorphin (5 μM), YAP signaling inhibitor verteporfin (100 nM), NOTCH signaling inhibitor DBZ (2 μM), SHH signaling inhibitor GDC-0449 (1 μM), WNT signaling inhibitor IWR-1 (5 μM), WNT signaling activator CHIR99021 (2 μM) which inhibits glycogen synthase kinase (GSK) 3, IL-6 (10 ng/ml), IL-4 (10 ng/ml) and IL-13 (10 ng/ml) were also used to treat ALI culture of EBCs.

### Treatment of mice with the Porcupine inhibitor Wnt-C59 and the γ secretase inhibitor DBZ

Wnt-C59 was resuspended by sonication for 20 minutes in a mixture of 0.5% methylcellulose and 0.1% tween-80. Wnt-C59 (10 mg/kg body weight) or vehicle was administrated via oral gavage from day 14 to 29 post viral infection (n=4 for each group). For DBZ administration, either vehicle or DBZ was administered intranasally at 30 mmol/kg body weight (n=4 per group) from day 14 to 29 post viral infection. DBZ was suspended in sterile PBS mixed with 2.5 μg/g body weight dexamethasone.

### Tissue and ALI culture processing and immunostaining

Human and mouse lung tissues were fixed in 4% paraformaldehyde (PFA) at 4°C overnight and then dehydrated and embedded in paraffin for sections (5-7 μm). ALI culture membranes were fixed with 4% PFA for direct wholemount staining or were embedded in OCT for frozen sections. All slides and membranes were stained following the protocol reported previously (Feldman et al., 2019; Mou et al., 2016). The primary antibodies used for immunostaining are listed in the Resources Table.

### Pulmonary function assessment

*Trpm5^+/-^* and *Trpm5^-/-^* mice (21 days after viral injury) were anesthetized with pentobarbital (50 mg/kg, i.p.). After surgical anesthesia was achieved, a tracheotomy was performed for the insertion of an 18G cannula, and mice were immediately connected to a flexiVent (SciReq, Montreal, QC, Canada) with an FX1 module and an in-line nebulizer. Body temperature was monitored and maintained with a thermo-coupled warming blanket and rectal temperature probe. Heart rate was monitored by EKG (electrocardiography). Mice were mechanically ventilated with a tidal volume of 10 mg/kg, frequency of 150 breaths/min, and a positive end expiratory pressure of 3 mmHg. Muscle paralysis was achieved with succinylcholine (10 mg/kg, i.p.) to prevent respiratory effort and to eliminate any contribution of chest wall muscle tone to respiratory measurements. By using the forced oscillation technique, baseline measurements of lung resistance (Rrs and Rn, representing total and central airway resistance) were performed. Resistance was then measured during nebulization of increasing concentrations of methacholine (10 second nebulization, 50% duty cycle). Methacholine dissolved in PBS was nebulized at 0, 6.25, 12.5, 25 and 50 mg/ml and resistance (Rrs and Rn) was measured after each concentration. Values for all measurements represent an average of triplicate measurements. Statistical significance was established by comparing the area under the curve for each mouse.

### COVID-19 lung specimens

The lung specimens from deceased COVID-19 patients with short post-mortem interval (PMI) (2.5-9hrs) were obtained from the Biobank at Columbia University Irving Medical Center. All experiments involving human samples were performed in accordance with the protocols approved by the Institutional Review Boards at Columbia University Irving Medical Center.

### Microscopic imaging and quantification

Slides were visualized using a ZeissLSM T-PMT confocal laser-scanning microscope (Carl Zeiss). The staining of cells on culture dishes and the staining on transwell membranes were visualized with the Olympus IX81 inverted fluorescence microscope. For quantification of lineage tracing, the lung sections were tiled scanned with 20X images from at least three mice for each genotype. Cells were counted from at least five sections per mouse including at least three individual lung lobes. The production of various airway epithelial cell types was counted and quantified on at least 5 random fields of view with a 10X or a 20X objective, and the average and standard deviation was calculated.

### Quantification and statistical analysis

Data are presented as means with standard deviations of measurements unless stated otherwise (n≥3). Statistical differences between samples are assessed with Student two-tailed T-test. P-values below 0.05 are considered significant (\**p*<0.05, \*\**p*<0.01, \*\*\**p*<0.001).

## Acknowledgements

We thank the colleagues in the Que laboratory for critical input of the study. We also thank Columbia University COVID-19 Hub and the autopsy pathologists who have been fighting on the frontlines and provided us short post-mortem interval (PMI) specimens.

## Author Contributions

H.H. and J.Q. designed the study and analyzed the data. H.M. and J.Q. supervised the research. H.H. performed mouse injuries, lineage tracing, immunostaining, in situ, imaging, ALI culture and mouse genetics. M.J. and Y.Z performed ALI culture. J.A.D. performed lung functional assays. M.J. and Y.F. assisted with moues injury models. J.L. assisted with COVID-19 sample immunostaining. J.B., J.C.M., Y.Y, L.Q., Y.Z., M.W., Z.H., T.C.W., A.S., J.S., I. M., W.C., C.W.E., J.Z., and B.I. assisted with data analysis. T.C.W. provided the *Trpm5^-/-^* and *Dclk1-GFP* mouse lines. H.H., J.Q. and H.M. wrote the manuscript with input from all authors.

## Data and materials availability

All data are available in the main text or the supplementary materials.

---

**Supplement file 1.**
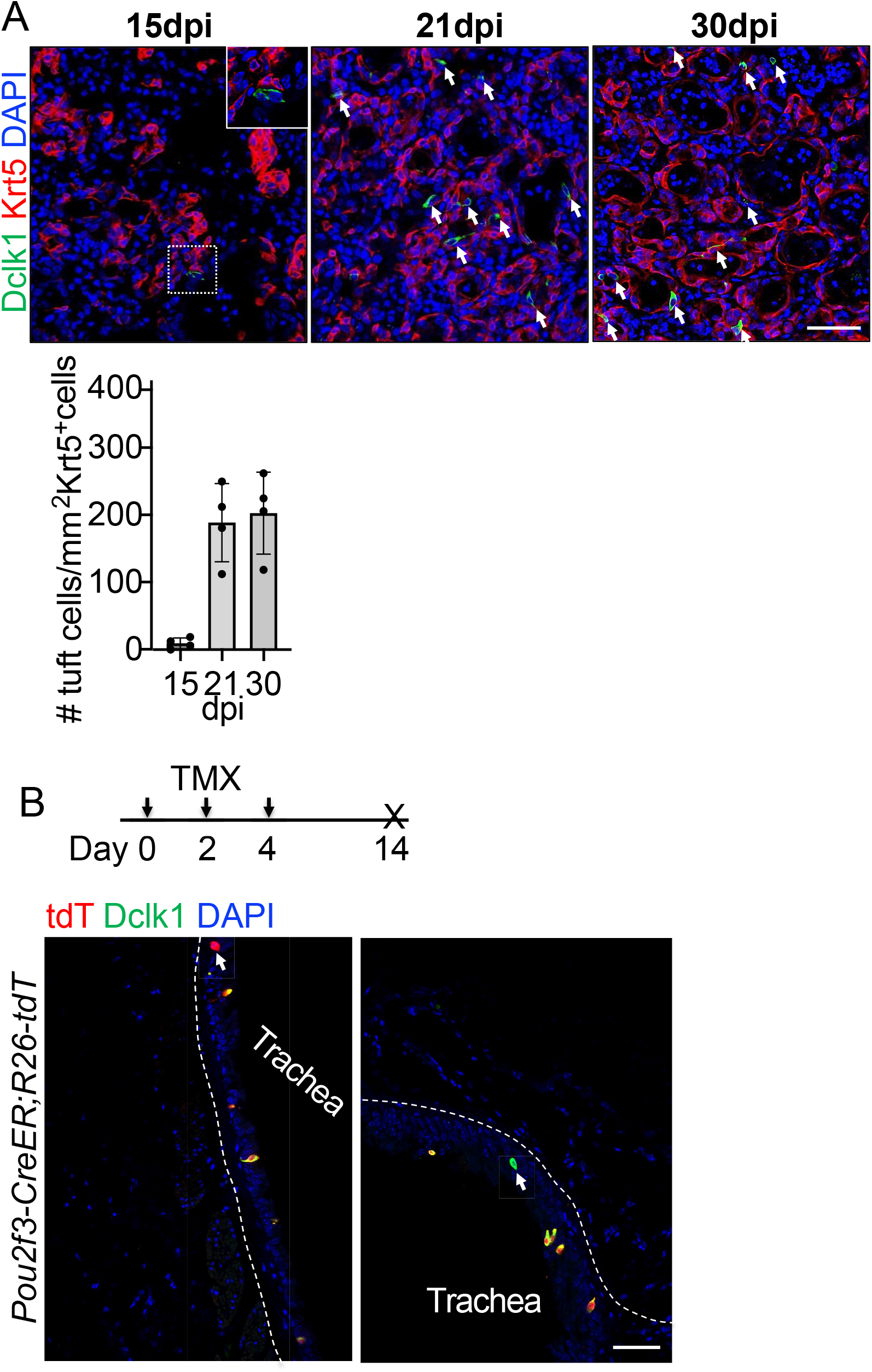
Tuft cells do not contribute to other cell lineages. (*A*) Time course analysis of ectopic tuft cells in the parenchyma following viral infection. (*B*) Lineage tracing shows heterogeneity of tuft cells in the trachea of *Pou2f3-CreER;R26-tdT* mice. Note that some lineage labeled cells do not express Dclk1 (arrowhead in left panel). Scale bars, 50 μm.

**Supplement file 2.**
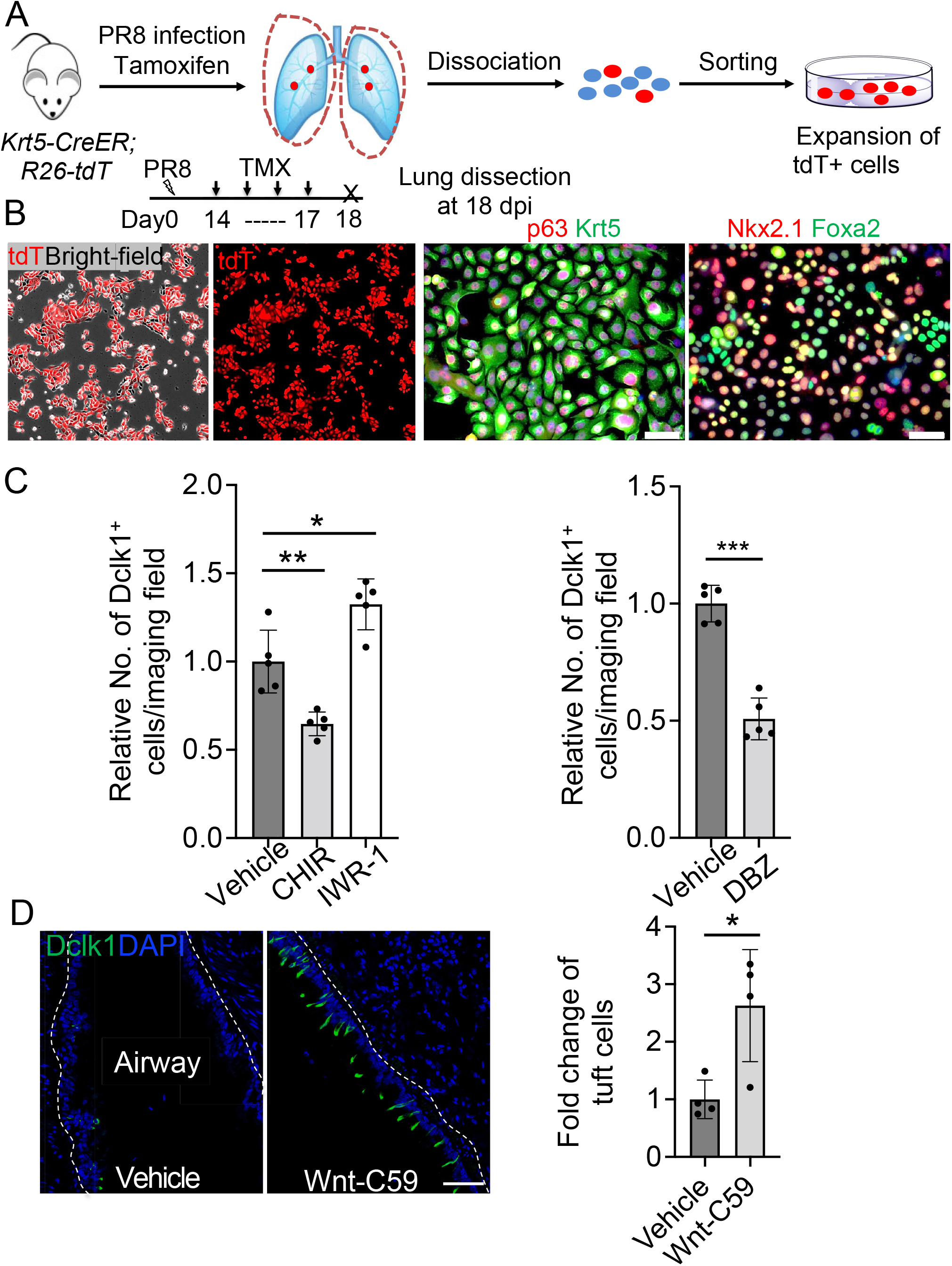
Isolation and expansion of Krt5^+^ EBCs from the lungs of *p63-CreER; R26tdT* mice that were infected with PR8 virus. (*A*) Schematic illustration depicts the methodology. (*B*) Lineage labeled EBCs are expanded and characterized. (*C*) Quantification of Dclk1^+^ cells derived from EBCs at various conditions (Vehicle, 2 μM CHIR99021, 5 μM IWR-1, and 2 μM DBZ) in air-liquid interface (ALI) culture. (*D*) Wnt-C59 treatment increases the numbers of tuft cells in the airways following viral infection. Data represent mean ± s.e.m. \**p*<0.05, \*\**p*<0.01, \*\*\**p*<0.001; statistical analysis by unpaired two-tailed Student’s *t*-test. Scale bars, 50 μm.

**Supplement file 3.**
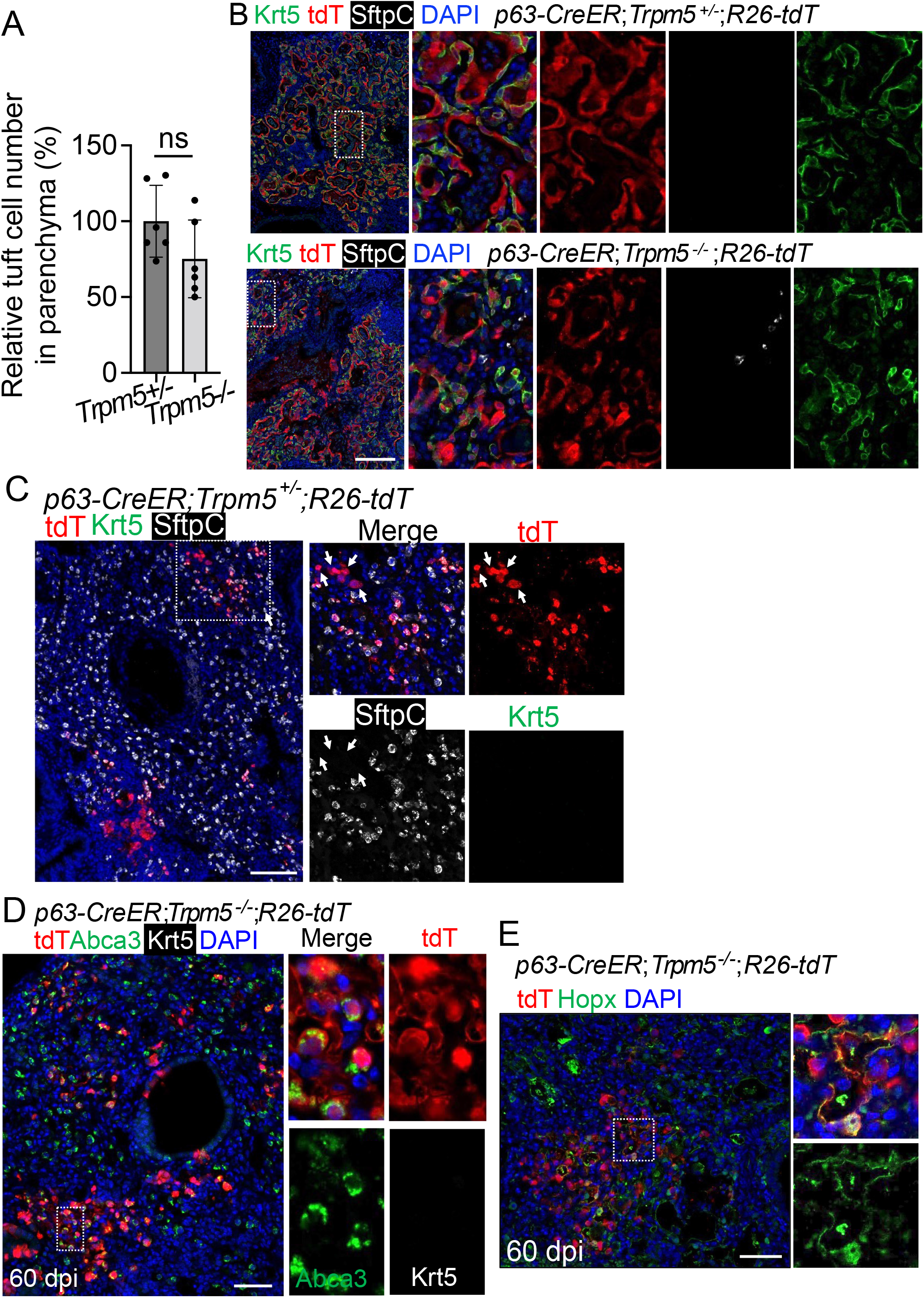
Increased *p63-creER* labeled AT1 and AT2 cells in virus-infected *Trpm5^-/-^* mice. (*A*) Quantification of tuft cells within EBCs established in the parenchyma of *Trpm5^+/-^* and *Trpm5^-/-^* mutants at 30 dpi. (*B*) Extensive Krt5^+^ EBCs in the lung parenchyma of controls (*p63-CreER;Trpm5^+/-^;R26tdT*) and mutants (*p63-CreER;Trpm5^-/-^;R26tdT*) at 60dpi. (*C*) Clusters of lineage-labeled AT2 cells. Note some lineage-labeled cells are Sftpc^-^ (arrowheads). Also note the same image in Figure4A. (*D*) Staining with Abca3 confirms the presence of lineage labeled AT2 cells in *p63-CreER;R26tdT;Trpm5^-/-^* mutants at 60 dpi. (*E*) Lineage tracing confirms that the presence of lineage labeled AT1 (Hopx^+^) in the parenchyma of *p63-CreER;R26tdT;Trpm5^-/-^* mutants. Data represent mean ± s.e.m. ns: not significant; statistical analysis by unpaired two-tailed Student’s *t*-test. Scale bars, 100 μm (B), 50 μm (C and D) and 20 μm (E).

Resources Table.

Resources Table
| Reagent or Resource | Source | Identifier |
| --- | --- | --- |
| **Primary Antibodies** |  |  |
| Dclk1 | Abcam | ab37994 |
| Scgb1a1 (mouse) | Santa Cruz | sc-365992 |
| SCGB1A1 (human) | R&D | MAB4218 |
| Krt5 | BioLegend | 905901 |
| Krt5 | Abcam | ab17130 |
| Krt5 | Abcam | ab53121 |
| p63 | Genetex | GTX102425 |
| p63 | Abcam | ab735 |
| Acetylated tubulin | Sigma | T7451 |
| Nkx2.1 | Abcam | ab76013 |
| Foxa2 | Abcam | ab60721 |
| Pou2f3 | Sigma | HPA019652 |
| Abca3 | Seven Hills | WRAB-70565 |
| Hopx | Santa Cruz | sc-398703 |
| SftpC | Abcam | ab90716 |
| Pro-SPC | Seven Hills | WRA3-9337 |
| Tomato | Biorbyt | orb182397 |
| T1α | DSHB | 8.1.1-c |
| GFP | Fisher (Aves Lab) | GFP1020 |
| Ki67 | Invitrogen | 14-5698-82 |
| **Secondary Antibodies** |  |  |
| Alexa Fluor 488 Donkey Anti-Chicken | Jackson Immuno Research | 703-546-155 |
| Alexa Fluor 488 Donkey Anti-Mouse | Jackson Immuno Research | 715-545-151 |
| Cy<sup>TM</sup>5 AffiniPure Donkey Anti-Rat IgG | Jackson Immuno Research | 712-175-150 |
| Cy<sup>TM</sup>3 AffiniPure Donkey Anti-Mouse IgM | Jackson Immuno Research | 715-165-140 |
| Alexa Fluor 488 Donkey anti-Rat IgG | Invitrogen | A21208 |
| Alexa Fluor 555 Donkey anti-Rabbit IgG | Invitrogen | A31572 |
| Alexa Fluor 555 Donkey anti-Goat IgG | Invitrogen | A21432 |
| Alexa Fluor 555 Donkey anti-Mouse IgG | Invitrogen | A31570 |
| Alexa Fluor 647 Goat anti-Chicken IgG | Invitrogen | A21449 |
| **Other reagents** |  |  |
| DBZ | Tocris | 4489 |
| Wnt-C59 | Tocris | 5148 |
| CHIR99021 | Tocris | 4423 |
| IWR-1 | Tocris | 3532 |
| Dorsomorphin | Sigma | P5499 |
| Verteporfin | Sigma | SML0534 |
| GDC-0449 | Selleckchem | S1082 |
| Dexamethasone | Sigma | D2915 |
| 3-Isobutyl-1-methylxanthine (IBMX) | Sigma | I5879 |
| Recombinant Il-6 | Peprotech | 200-06 |
| Recombinant Il-4 | Peprotech | 200-13 |
| Recombinant Murine Il-13 | Peprotech | 210-13 |
| Normal donkey serum | Jackson Immuno Research | 017-000-121 |
| 8-Bromoadenosine 3',5'-cyclic<br>monophosphate (8-Br-cAMP) | Sigma | B5386 |
| Bleomycin | Fresenius Kabi | 63323013610 |
| Naphthalene | Sigma-Aldrich | 84679 |
| Antigen unmasking solution, Citric acid<br>based | Vector Laboratories | H-3300 |
| Small Airway Epithelial Cell Medium | Lonza | CC-3118 |
| Small Airway Epithelial Cell Medium | Promocell medium | C-21170 |
| Complete Pneumacult-ALI medium | StemCell Technology | 05001 |

